# The NHEJ Repair of DNA Double Strand Breaks in *Physcomitrella patens* Depends on the Kleisin NSE4 of the SMC5/6 Complex

**DOI:** 10.1101/2020.05.21.108175

**Authors:** Radka Vágnerová, Marcela Holá, Karel J. Angelis

## Abstract

Structural maintenance of chromosomes (SMC) complexes are involve in cohesion, condensation and maintenance of genome stability. Based on the sensitivity of mutants to genotoxic stress the SMC5/6 complex is thought to play imminent role in DNA stabilization during repair by encircling DNA at the site of lesion by bridging the heteroduplex of SMC5 and SMC6 by non SMC kleisin components NSE1, 3 and 4. In this study, we tested how formation of the SMC5/6 circular structure affects mutant sensitivity to genotoxic stress, kinetics of DSB repair and insertion mutagenesis. In the moss *Physcomitrella patens SMC6* and *NSE4* are essential single copy genes and this is why we used blocking of transcription to reveal their mutated phenotype. Even slight attenuation of transcription by dCas9 binding was enough to obtain stable lines with DSB repair defect and specific bleomycin sensitivity. Whereas survival after bleomycin or MMS treatment fully depends on active SMC6, NSE4 has little or negligible effect. We conclude that whereas circularization of SMC5/6 provided by the kleisin NSE4 is indispensable for the immediate NHEJ DSB repair response, other functions associated with SMC5/6 complex are critical to survive DNA damage.

## INTRODUCTION

Plants as sessile organisms developed several strategies to protect integrity of their genomes against various environmental issues including genotoxic stress. The bryophyte *Physcomitrella patens* (*P. patens*) stands aside from other plants with high frequency of homologous recombination (HR) due to its remarkable ability to integrate transgenes at predefined loci through homologous recombination-mediated gene targeting (GT) (Kamisugi et al. 2016). Experimentally, *P. patens* is highly amenable to analysis and manipulation e.g. for study of DNA damage and repair by comet assay (Angelis et al. 2015). *P. patens* spore develops into a haploid filamentous structure called protonema, composed of two types of cells – chloronema and caulonema. With respect to *in vitro* cultivation conditions in basal medium enriched with ammonium tartrate (Knight et al. 2002) protonema is exclusively composed of chloronemal cells accumulated at the G2/M transition. This unique tissue-specific cell cycle arrest is thought to be behind uniquely high rate of HR in *P. patens* nuclear DNA (Schween et al. 2003). Protonema filaments grow exclusively by tip growth of their apical cells (Menand et al. 2007). The advantage of using filamentous *P. patens* is that it can be employed either as a culture of dividing cells approximated by 3-7 cell fragments with average 50% of apical cells obtained from protonemal lawns by extensive shearing or as differentiated tissue from 7-days grown protonemata with only 5% of apical cells. Either culture exhibits different kinetics of DNA repair (Goffova et al. 2019; Kamisugi et al. 2012). APT (<u>a</u>denine <u>p</u>hosphoribosyltransferase) is an enzyme of the purine salvage pathway that converts adenine into AMP and its loss of function generates plants resistant to adenine analogues e.g. 2-fluoroadenine (2-FA) (Gaillard et al. 1998). Mutational inactivation can be used as selectable marker as well as analysis of mutations in *APT l*ocus on nucleotide level (Trouiller et al. 2007; Trouiller et al. 2006).

The SMC6 (<u>s</u>tructural <u>m</u>aintenance of <u>c</u>hromosome) protein is component of highly conserved SMC5/6 complex that is composed of SMC5 and SMC6 heterodimer and 6 non-SMC elements NSE1-6. Similarly to the other SMC complexes of cohesin and condensin, SMC5-SMC6 heterodimer together with NSE4 klesin-bridge form circular structure capable of entrapping DNA (Hassler et al. 2018). The NSE1 and NSE3 subunits regulate directly or indirectly the ring shape and functions (Vondrova et al. 2019). This conserved core of the SMC5/6 complex is indispensable for all its functions including essential processes like DNA replication and repair (Diaz and Pecinka 2018; Palecek 2018). *NSE4* and *SMC6* are essential genes in yeast and mammals and their knockout is lethal (Hu et al. 2005; Ju et al. 2013; Onoda et al. 2004). The *Arabidopsis* orthologue *AtSMC6b* (*MIM*) was shown to interfere with DSBs repair by eliminating its rapid phase (Kozak et al. 2009). Because the haploid state of the *P. patens* protonema and indispensability of the SMC5/6 complex do not allow complete depletion of any of the complex subunits we used attenuated expression approach by specific binding of catalytically dead Cas9 (dCas9) without endonucleolytic activity to desired gene and generate mutants with attenuated transcription. Such mutant plants are still viable, growing and manifesting DNA repair phenotype.

## MATERIALS AND METHODS

### Plant Material and Cultivation

The WT *Physcomitrella patens* (accession (Hedw.) B.S.G. ‘Gransden2004’) (Knight et al. 2002), the *ppku70* and *pprad51-1-2* double mutant (Schaefer et al. 2010) were generated by D. G. Schaefer, Neuchatel University, Switzerland, and F. Nogué, INRA, Paris, France, and kindly provided by F. Nogué. All strains were cultured as ‘spot inocula’ on BCD agar medium supplemented with 1mM CaCl2 and 5mM ammonium tartrate (BCDAT medium), or as lawns of protonema filaments by subculture of homogenized tissue on BCDAT agar overlaid with cellophane in growth chambers with 18/6 h day/night cycle at 22/18**°**C (Knight et al. 2002).

One-day-old protonema tissue (1d) for repair experiments was prepared from 1-week-old tissue (7d) scraped from plate, suspended in 8 ml of BCDAT medium, sheared at 10 000 rpm for two, 1-min cycles by a T25 homogenizer (IKA, Germany) and let to recover for 24 h in cultivation chamber with gentle shaking at 100 rpm. This treatment yielded a suspension of 3**–**5 cell protonemata filaments, which readily settle for recovery.

### Generation and Analysis of *PpNSE4* and *PpSMC6* Mutants

The vectors for attenuation of *PpNSE4* and *PpSMC6* expression were constructed as expression cassette of dCas9 driven by maize ubiquitin promoter, NPTII selection cassette and specific guide RNA (sgRNA). Construction, transformation and validation of mutated lines is described in Supplementary Methods S1.

Three stable transformants of *ppnse4* and *ppsmc6* were randomly picked and analyzed by quantitative RT-PCR (qRT-PCR) for *NSE4* or *SMC6* expression (Fig. 1b). Total RNA was isolated from 7 days old protonemata with RNeasy Plant Mini Kit (Qiagen), treated with DNaseI (DNA-*free*™ DNA Removal Kit, Thermo Fisher Scientific) and reverse transcribed using qPCRBIO cDNA Synthesis Kit (PCR Biosystems). Diluted cDNA reaction mixtures were used for qRT-PCR analysis using the qPCRBIO SyGreen Mix Lo-ROX (PCR Biosystems) in Statagene-MX3005P. Analysis was performed for three biological replicas (independently cultivated tissue) and in two technical replicates with Clathrin adapter complex subunit CAP-50 (Pp3c27_2250V3.1) as a reference gene (Kamisugi et al. 2016). The relative transcription of *NSE4* and *SMC6* was calculated by the ΔΔCt method (Pfaffl 2004).

**FIG. 1.**
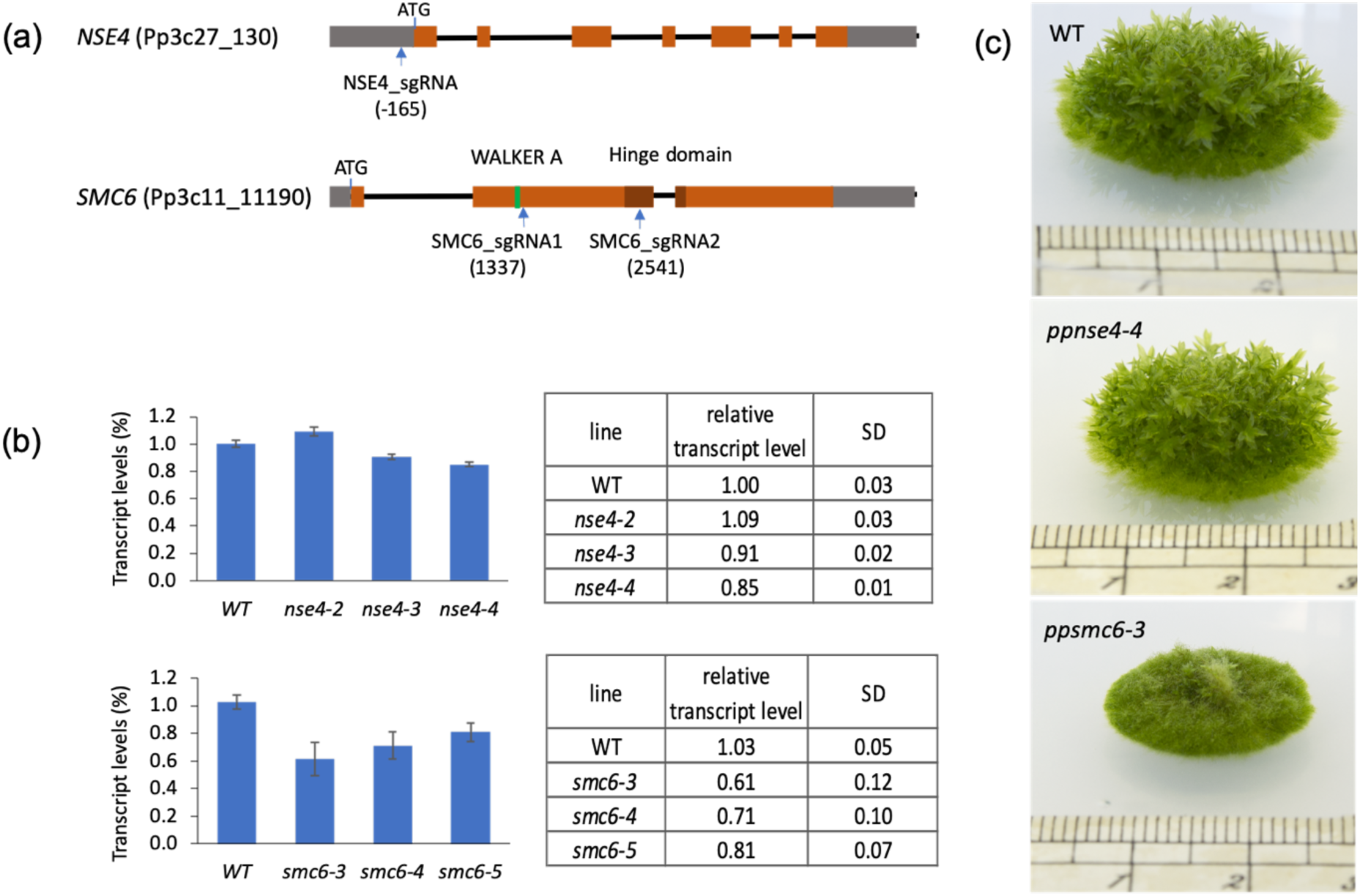
Characterization of *ppnse4* and *ppsmc6* mutants. (a) Structure of the WT, PpNSE4, and PpSMC6 loci with indication of dCas9 binding. The brown boxes represent exons and the gray’s UTR regions. (b) qRT-PCR analysis of *NSE4* (upper) and *SMC6* (lower) transcripts in WT and mutant plants. Primers are listed in Supplementary Table S1. (c) Morphology of colonies of WT and of *ppnes4-4* and *ppsmc6-3* mutants. The picture was taken after 1.5 month of growth.

### Mutagens, Treatments and Sensitivity Assay

Sensitivity to DNA damage was measured after treatment either with freshly prepared solutions of bleomycin sulphate supplied as Bleomedac inj. (Medac, Hamburg, Germany), or methyl methanesulfonate (MMS) (Sigma-Aldrich) in BCDAT medium. Protonemata were dispersed in liquid BCDAT medium containing bleomycin or MMS and treated for 1 h. After treatment and washing recovered protonemata were inoculated as eight explants per quadrant of a Petri dish with drug-free BCDAT agar without cellophane overlay and left to grow. UVC irradiation was performed in a Hoefer UVC 500 crosslinker at 254 nm by irradiating *P. patens* lines spotted as explants on a Petri dish. Irradiation and the following steps were performed for 24 h in the dark or under red illumination to block photolyases. Treatment was assessed after 3 weeks by weighing explants. The fresh weight of the treated explants was normalized to the fresh weight of untreated explants of the same line and plotted as % of ‘Relative fresh weight’. In every experiment, the treatment was carried in duplicate and experiments were repeated two or three times and statistically analyzed by Student t-test.

### Single-Cell Gel Electrophoresis (Comet) Assay

Repair kinetics was estimated in 1d and 7d protonemata after bleomycin or MMS treatment. Tissue was either flash-frozen in liquid N2 (repair t **=** 0) or left to recover in liquid BCDAT medium for the indicated repair times and then frozen. DSBs after bleomycin treatment were detected by a comet assay using neutral N/N protocol whereas SSBs after MMS treatment were detected with A/N protocol, which includes after lysis of nuclei a treatment with alkali to reveal breaks by unwinding DNA double helix as described in (Angelis et al. 1999; Hola et al. 2013). Comets were stained with SYBR Gold (Molecular Probes/Invitrogen) and viewed in epifluorescence with a Nikon Eclipse 800 microscope and evaluated by the LUCIA Comet cytogenetic software (LIM Inc., Prague, Czech Republic).

The fraction of DNA in comet tails (% tail-DNA) was used as a measure of DNA damage. In each experiment, the % tail-DNA was measured at seven time points: 0, 3, 5, 10, 20, 60 and 180 min after the treatment and in control tissue without treatment. Measurements obtained in at least three independent experiments and totaling at least 300 comets analyzed per experimental point were plotted as % of remaining damage and statistically analyzed by Student t-test.

### Gene Targeting Assay

GT efficiencies were assessed after transformation with the APT based targeting construct pKA255 (Fig. 4). Vector pKA225 contains 35S-Hygromycin marker (HygroR) flanked by 775bps 5’-homology fragment of *APT* locus (Pp3c8_16590V3.1) (position −766 - 9 across the start codon) and 622 bps 3’-homology fragment (658 – 1280) that were amplified from moss genomic DNA by PCR. Linear targeting fragment was amplified by PCR and delivered by PEG mediated transformation to WT and *ppnse4-4* and *ppsmc6-3* protoplasts (Liu and Vidali 2011). Transformed protoplasts of each line were spread on four Petri dishes with BCDAT medium overlaid with cellophane disc. Regenerating protoplasts was counted after 5 days on each Petri dish and then transferred to new Petri dish with BCDAT medium supplemented with 5 μM 2-FA. Successful transformation was followed by direct counting of 2-FA resistant (2-FA^R^) colonies after 3 weeks. Frequency of GT is expressed as the ratio between the number of 2-FA^R^ colonies and the number of regenerating protoplasts. Experiments were repeated three times and statistically analyzed using the Fisher exact test.

**FIG. 2.**
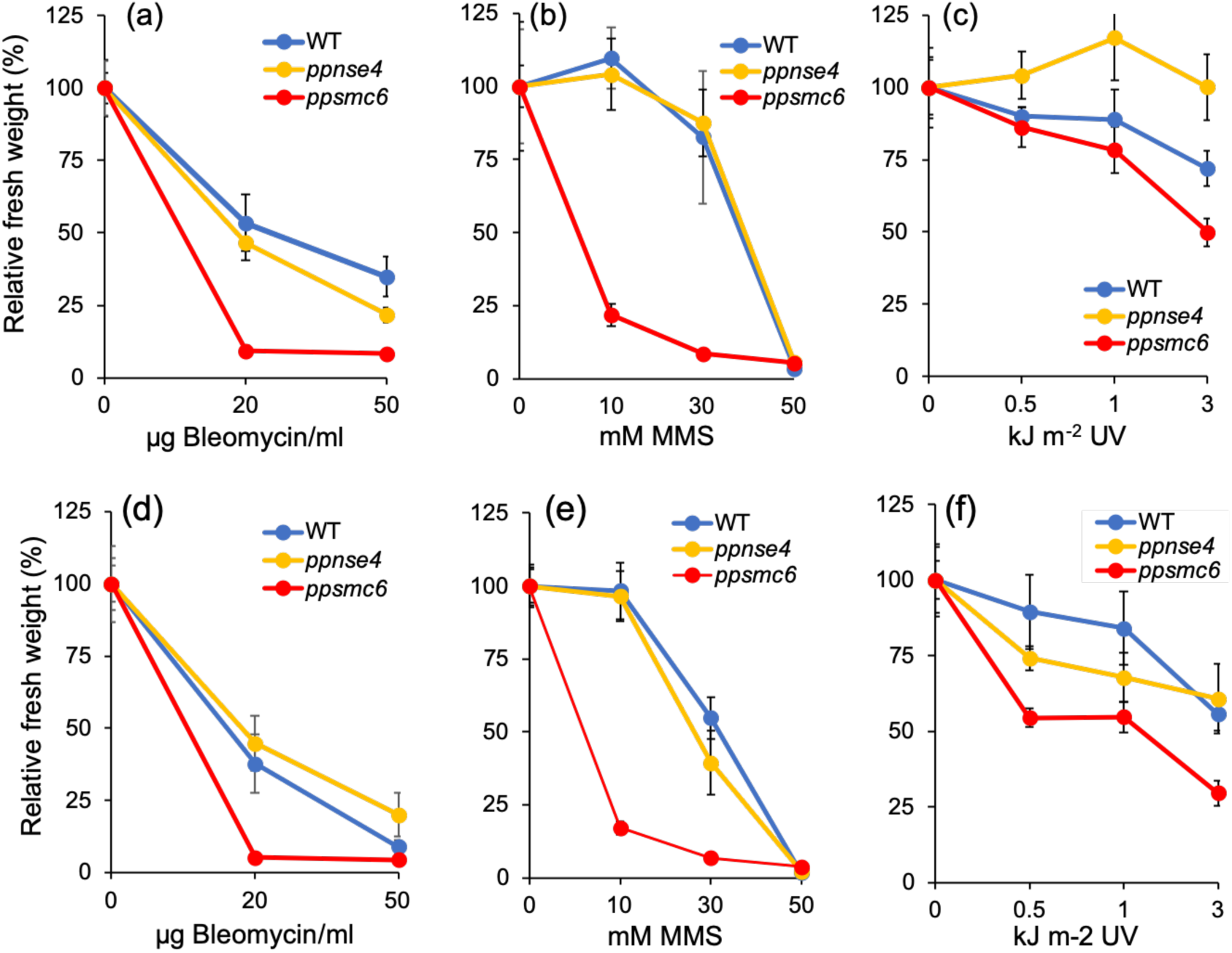
Growth-responses of 1d (a,b,c) and 7d (d,e,f) protonemata of *P. patens* WT, *nse4-4* and *smc6-3* to 1h treatment with bleomycin (a, d), MMS (b, e) or UVC-irradiation (c, f). After treatment, the explants were incubated on drug-free BCDAT medium under standard growth conditions for 3 weeks. For each experimental point the weight of treated plants collected from two replicas was normalized to the weight of untreated plants and plotted as relative fresh weight, which was set as a default to 100. Error bars indicate SD.

**FIG. 3.**
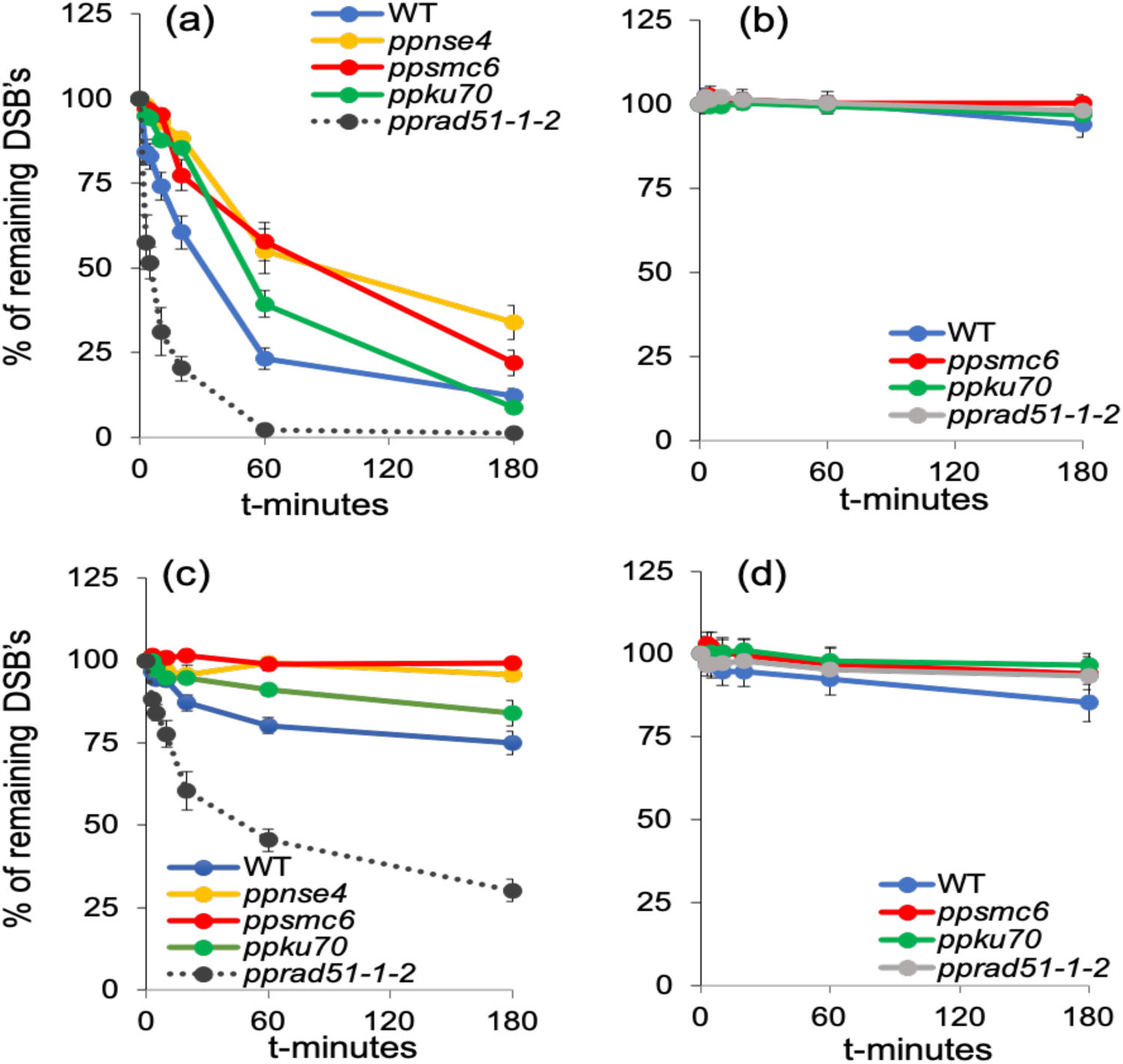
DSB and SSB repair kinetics determined by comet assay in 1d or 7d tissue of WT (blue) and *ppnse4,4* (orange), *ppsmc6-3* (red), *ppku70* (green) and *pprad51-1-2* (gray) mutants. Data for *pprad51-1-2* (a,c) are from (Goffova et al. 2019). Protonemata regenerated 1d (a,b) or 7d (c,d) after subculture were treated with 30 µg bleomycin/ml or 30 mM MMS for 1h and repair kinetics was measured as % of DSBs (a,c) or SSB (b,d) remaining after the 0, 3, 5, 10, 20, 60 and 180 min of repair recovery. Maximum damage is normalized as 100% at t=0 for all lines. Error bars indicate SD.

**FIG. 4.**
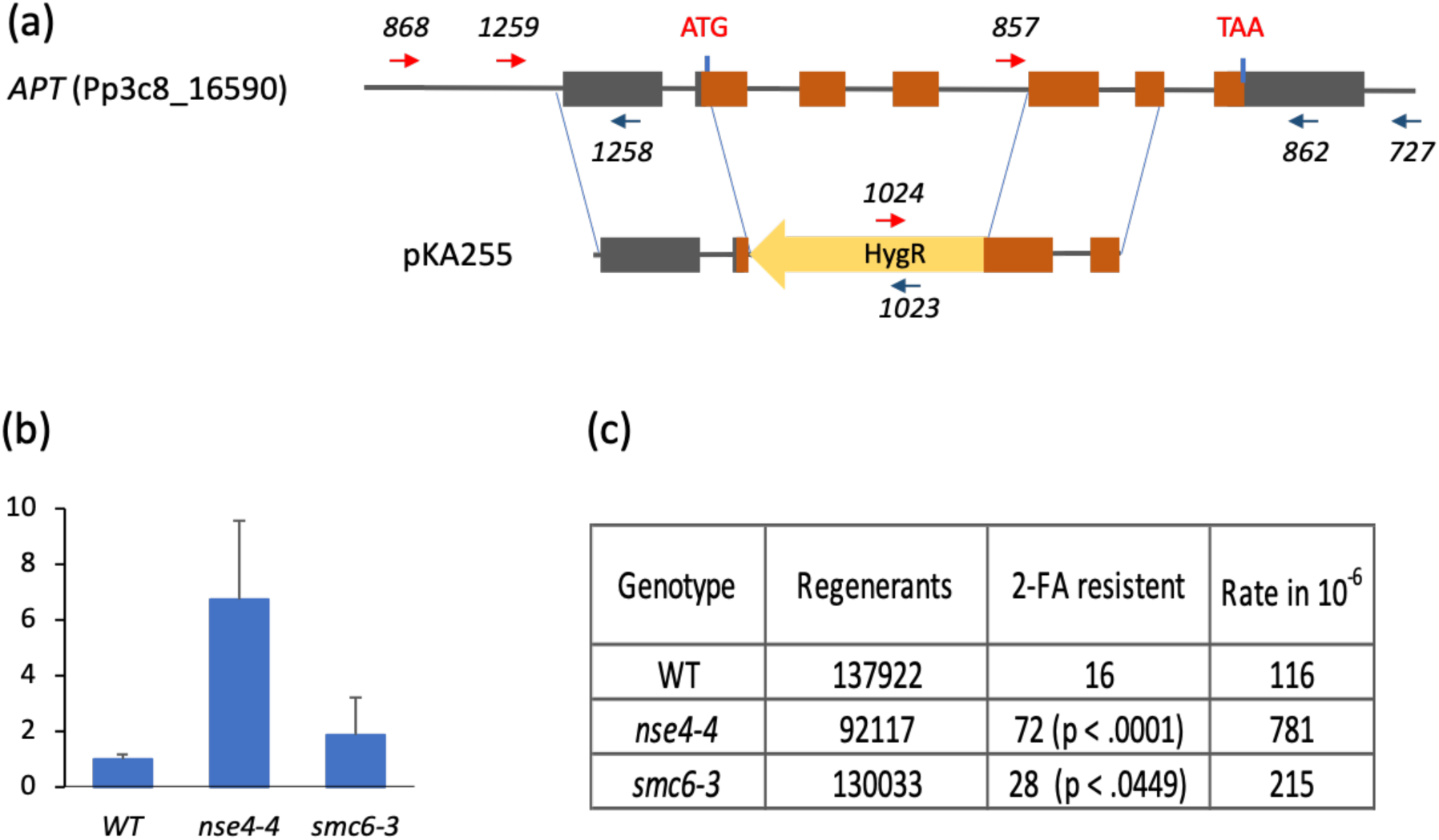
GT assay. (a) Schematic drawing of *APT* locus and of targeting construct pKA255. The brown boxes represent exons and the gray’s UTR regions. Arrows mark position of primers used to genotype and sequence the plants by PCR. (b) GT efficiency in *ppnse4-4* and *ppsmc6-3* in comparison to WT, which was set as a default to 1). (c) The rate of GT per 10^6^ protoplasts. Differences between WT and mutants were compared using Fisher’s exact test.

In recovered 2-FA^R^ clones integration of pKA255 was checked by PCR in six randomly picked 2-FA^R^ clones of each line with external gene primers *868* and *727* and selection cassette primers *1023* and *1024* and integration borders were confirmed by sequencing with primers *862, 857, 1259* and *1258* (Supplementary Table S1).

### Statistical analysis

Three biological repeats were conducted for each experiment. Statistical significance between two samples was analyzed by two-tailed Student’s t test.

## RESULTS and DISCUSSION

### Generation and Analysis of *ppnse4* and *ppsmc6* Mutants

Based on protein sequence of *A. thaliana* search in *P. patens* genome (Phytozome v12.1) we identified single copy locus of *NSE4* (Pp3c27_130V3.1) and of *SMC6* (Pp3c11_11190V3.1). In *P. patens NSE4* and *SMC6* are essential genes and their deletion is lethal (unpublished observation), nevertheless it is possible to obtain stable dCas9_sgRNA transformants with targeted *NSE4* promotor and *SMC6* hinge region (Fig. 1a). It was not possible to recover any stable dCas9 transformant with targeted *SMC6* promotor or ATPase Walker A region.

Three rescued lines of *NSE4*^-^ (*ppnse4-4,5* and *6*) and of *SMC6*^-^ (*ppsmc6-1,2,3*) were analyzed for remaining expression by qRT-PCR (Fig. 1b). In this study we use *ppnse4-4* with 15% an*d ppsmc6-3* with 40% reduction of transcription, both the highest values in rescued lines. Phenotype of *ppnse4-4* though growing slightly slower do not manifest any obvious deviation from WT, whereas *ppsmc6-3* growth is reduced, and plants do not develop gametophores (Fig. 1c).

The attenuation was repeatedly confirmed in both lines after several months.

### Recovery from Bleomycin and MMS Treatment Depends on PpSMC6

To characterize defects in *ppnse4-4, smc6-3, ppku70* and *pprad51-1-2* mutants we compared their growth-responses to lesions like DSBs induced by radiomimetic bleomycin, small alkylation adducts induced by MMS and DNA helix distortion by photo adducts induced by UVC irradiation. These lesions represent blocks for DNA replication and are removed, repair or bypassed by various error-free as well as error-prone pathways. The sensitivity was tested after acute treatment for 1h as the ability of the tissue to recover.

In response to the treatment with bleomycin or MMS, sensitivities of *ppnse4-4* and *ppsmc6-3* substantially differed, both in cultures with dividing (1d) as well as differentiated (7d) cells. Whereas the sensitivity of *ppnse4-4* was very similar to WT, *ppsmc6-3* was extremely vulnerable (Fig. 2a,b,d,e). After exposure to 20 µg bleomycin/ml and 30 mM MMS *ppsmc6-3* explants did not recover at all, while WT and *ppnse4-4* still retain about 50%, and in 1d culture treated with 30 mM MMS even 100% viability (Fig. 2b).

The response to bleomycin and MMS suggests that formation of circular form of SMC5/6 complex is not inevitably necessary to offset toxic effect of DNA lesions and so enable cell survival, but availability of complex itself and additional associated activities like SUMOylation and other so far undisclosed functions are critical and essential for survival. Response to UVC-irradiation does not give clear picture on the role of *NSE4* and *SMC6* in the response to induced lesions, nevertheless *ppsmc6-3* remain the most vulnerable in dividing as well as differentiated tissue.

### DSB Repair is Impaired in *nse4* and *smc6*

DSB repair is severely inhibited in both *ppnse4-4* and *ppsmc6-3* lines where both share identical repair kinetic in dividing (1d) as well as differentiated (7d) cells (Fig. 3a,c). In this respect they closely parallel previously reported repair defect in *A. thaliana* mutant *atmim* (*atsmc6b*) (Kozak et al. 2009), when SMC6b was suggested to participate in NHEJ due to absence of repair rapid phase that is assumed to represent NHEJ repair (Goodarzi et al. 2010). To ascertain the repair pathway where SMC6 is involved, we compared *ppnse4-4* and *ppsmc6-3* with cNHEJ mutant *ppku70* and HR mutant *pprad51-1-2*. Statistical analysis showed that actually their repair kinetic differs from both, *pprad51-1-2* (1d p < 0.01; 7d p < 0.05) and *ppku70* (1d and 7d p < 0.05). This result supports conclusion from *A. thaliana* that SMC5/6 is involved in tethering DNA strands during alternative NHEJ, distinct from cNHEJ also in *P. patens*. For tethering DNA and participation in DSB repair bridging of ATPase heads of SMC5/6 heteroduplex by kleisin NSE4 is critical what is documented by the fact that weak, 15%, attenuation of *NSE4* transcription results in the same extent of repair deficit as 40% attenuation of *SMC6* transcription.

MMS does not induce DSBs detectable by comet assay and repair of SSBs in WT and all mutants is completely abolished (Fig. 3b,d). This is in contrast with repair of bleomycin induced SSBs when in WT 50% of breaks are already removed after 3h (Hola et al. 2013).

### Gene Targeting is Increased in the *nse4* and *smc6* Mutants

The GT vector pKA225 was designed to disrupt *PpAPT* gene across the second exon site (229-339) previously identified as mutation hotspot (Hola et al. 2013; Hola et al. 2015). The GT rate was determined as the frequency of 2-FA^R^ plants amongst 10^6^ regenerating protoplasts (Fig. 4c). There is a statistically significant increase of GT rate after attenuation of *NSE4* and *SMC6*. As evident from Fig. 4b the rate of GT is increased approximately 7x (p < 0001) in *ppnse4-*4 and only 2x (p < 0.049) in *ppsmc6-3* over WT. PCR analysis of 6 randomly selected 2-FA^R^ plants of each line revealed correct, full length integration. To assess accuracy of integration, insert borders were sequenced in two 2-FA^R^ transformants of each line and all analyzed transformants had accurate insertion at both ends of insert without detectable mutation. This result supports above described finding that attenuation of *SMC6* and particularly of *NSE4* expression impairs error-prone NHEJ repair machinery of whatever mechanism is involved during integration and shift balance to error-free HR insertion.

## CONCLUSION

In this study we used haploid *P. patens* cultivated on rich BCDAT medium as chloronema with cells arrested at G2/M border to study the role of SMC5/6 complex in DNA repair. SMC5/6 in circular form is indispensable for DSB repair presumably by NHEJ, but not cNHEJ pathway, what is illustrated by the same repair kinetic of the *smc6* and kleisin *nse4* mutants that is distinct from *ku70*. Nevertheless, the repair dependent on NSE4 is not essential for cell survival after bleomycin and MMS treatment as the incurred damage can be offsetted by alternative repair mechanisms. The increased rate of GT shows that defect or elimination of NSE4 dependent pathway shifts equilibrium in favor of undisturbed HR mechanism. Still remain question, which functions and qualities of SMC5/6 enable survival of *nse4* after bleomycin or MMS treatment, but are disturbed in *smc6*, or whether the observed differences are just consequence of different degree of *NSE4* and *SMC6* expression.

## ACKNOWLEDGEMENTS

The research was supported by the Czech Science Foundation (project 20-05095S) and by the Ministry of Education, Youth and Sports of the Czech Republic under the project COST (LTC17047).

## AUTHOR CONTRIBUTIONS

MH and KJA designed the research, analyzed data and wrote the manuscript. RV, MH and KJA conducted the research.

## CONFLICT OF INTERESTS

The authors declare no conflicts of interest.

## SUPPLEMENTARY MATERIALS

The Supplementary Material for this article can be found online at:

## Notes

### Competing Interest Statement

The authors have declared no competing interest.

