## Supplementary Methods for "The NHEJ Repair of DNA Double Strand Breaks in *Physcomitrella patens* Depends on the Kleisin NSE4 of the SMC5/6 Complex"

### Methods S1 dCas9 attenuation of *PpNSE4* and *PpSMC6* expression

**(a)** The vectors for attenuation of *NSE4* and *SMC6* expression in *P.patens* were constructed as dCas9 expression cassette driven by maize ubiquitin promoter, NPTII selection cassette and specific guide RNA (sgRNA) under the *AtU6_26* promoter (Fauser et al., 2014) in Golden Braid vector pDGBΩ1(Figure S1)(Sarrion-Perdigones et al., 2013).

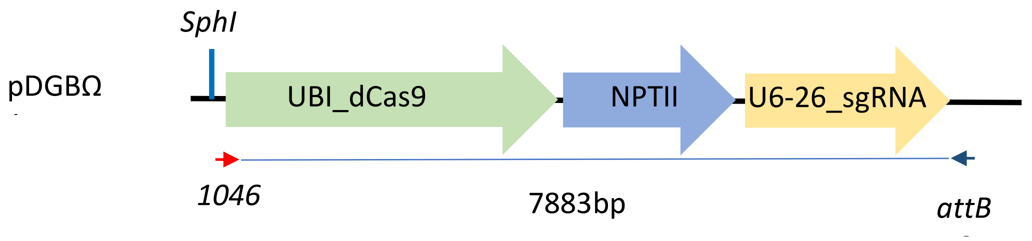

FIGURE S1 Schematic drawing of dCas9 attenuation vector

*NSE4* sgRNA targets binding of dCas9 to promotor at -165nt upstream of start codon. For *SMC6*, two sgRNA's were designed. SMC6_sgRNA1, which targets dCas9 closely to WALKER A motif of ATPase head at position 1338 and SMC6_sgRNA2 that targets position 2541nt at the sequence coding hinge domain (Figure S2).

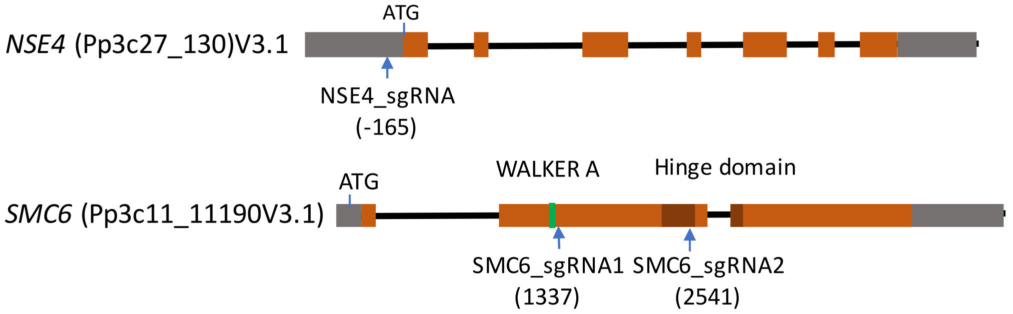

FIGURE S2 – Targeted positions in NSE4 and SMC6 indicated by blue arrows.

The sgRNA's were designed according coding sequences of *PpNSE4* (Pp3c27_130V3.1) and *PpSMC6* (Pp3c11_11190V3.1) with CRISPOR program (Haeussler et al., 2016) against *P. patens* genome Phytozome V9. sgRNAs fragments were obtained by annealing oligonucleotides 1092+1093 for *NSE4* and 1175+1176 for RNA1 and 1173+1174 for RNA2 of *SMC6*.

**(b)** *P. patens* protoplasts were transformed by PEG method (Liu and Vidali, 2011) with *SphI* linearized vectors and Stable transformants were obtained after three rounds of selection on medium containing 50 µg/ml G418.

**(c)** The integration of full-length constructs was verified by PCR (Figure S3) covering entire construct with primers *1046* and *attB2* flanking the construct. The PCR was performed from genomic DNA isolated from *ppnse4-4* and *ppsmc6*, respectively. Expected product size is 7883bps for both lines.

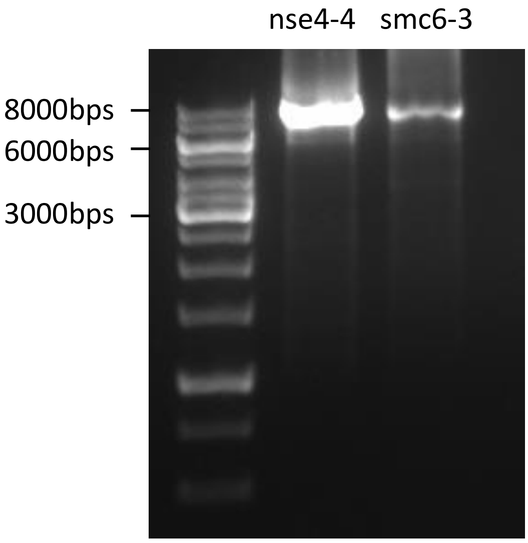

FIGURE S3 Integrated attenuation vectors in genome of *ppnse4-4* and *ppsmc6-3*

**(d)** Primers used:

| PCR primers | integrity of dCas9 construct | |
|---|---|---|
| 1046 | F | GCGCCGTCTCGCTCGGGAGGTCAGGAGCATGATTACGAAT |
| attB2 | R | GGGGACCACTTTGTACAAGAAAGCTGGGT |
| sgRNA oligos | sgRNA cloning | |
| 1173 | F | ATTGATCCAATCGGTCCAATGGG |
| 1174 | R | AAACCCCATTGGACCGATTGGAT |
| 1175 | F | ATTGCAGCGTGCCACGTCGCTGA |
| 1176 | R | AAACTCAGCGACGTGGCACGCTG |
| 1091 | F | ATTGTGTGGTTTCTTGATGACGG |
| 1092 | R | AAACCCGTCATCAAGAAACCACA |

### Table S1 Primers used in this study

| Sequencing primers | Purpose/orientation | |
|---|---|---|
| 857 | F | CAATGTGACCGAGACTTCATCC |
| 862 | R | CGTTTCCGAGGTTAGAAG |
| 1258 | R | TCACACCCTGAAGCCCTAGC |
| 1259 | F | TTAGAGCTATGACATTAGTG |
| PCR primers | integrity of dCas9 construct | |
| 1046 | F | GCGCCGTCTCGCTCGGGAGGTCAGGAGCATGATTACGAAT |
| attB2 | R | GGGGACCACTTTGTACAAGAAAGCTGGGT |
| PCR primers | pKA255 integration | |
| 727 | R | TCGCGGTTCAGATTCGACGG |
| 1024 | F | GATTCCTTGCGGTCCGAATG |
| 868 | F | TGACAAAGTGCTTCTATACC |
| 1023 | R | AGCGAGAGCCTGACCTATTG |
| qRT-PCR primers | qRT-PCR | |
| 839 | SMC6_R | TGCACTCTTTCCACTGCCATTC |
| 962 | SMC6_F | CTCCTGCGTTACATCAATCG |
| 1003 | NSE4_F | TGCTTCGCTTCGTTTCTGTC |
| 1004 | NSE4_R | TCGCTCTCCTGCTACTTATTCC |
| 1001 | CAP50_F | TACAATGCGGCTACCGAC |
| 1002 | CAP50_R | AGGCCGGATGCAGTAAAC |
| sgRNA oligos | protospacers cloning | |
| 1173 | F | ATTGATCCAATCGGTCCAATGGG |
| 1174 | R | AAACCCCATTGGACCGATTGGAT |
| 1175 | F | ATTGCAGCGTGCCACGTCGCTGA |
| 1176 | R | AAACTCAGCGACGTGGCACGCTG |
| 1091 | F | ATTGTGTGGTTTCTTGATGACGG |
| 1092 | R | AAACCCGTCATCAAGAAACCACA |
